# Convolutional neural network approach for the automated identification of *in cellulo* crystals

**DOI:** 10.1101/2023.03.28.533948

**Authors:** Amirhossein Kardoost, Robert Schönherr, Carsten Deiter, Lars Redecke, Kristina Lorenzen, Joachim Schulz, Iñaki de Diego

## Abstract

*In cellulo* crystallization is a rarely occurring event in nature. Recent advances, making use of heterologous overexpression, can promote the intracellular formation of protein crystals, but new tools are required to detect and to characterize these targets in the complex cell environment. In the present work we make use of Mask R-CNN, a Convolutional Neural Network (CNN) based instance segmentation method, for the identification of either single or multi-shaped crystals growing in living insect cells, using conventional bright field images. The algorithm can be rapidly adapted to recognize different targets, with the aim to extract relevant information to support a semi-automated screening pipeline, with the purpose to aid in the development of the intracellular protein crystallization approach.

## 1. Introduction

Crystallization of proteins in living cells is an emerging field complementing conventional methods of protein crystallization. However, some bottlenecks still limit the broad application. To date, around 80 targets have been found to crystallize in different cells (Tsukimoto et al., 2022; Li et al., 2020; Nass et al., 2020; Mudogo et al., 2020; Schönherr et al., 2018; Duszenko et al., 2015; Koopmann et al., 2012). Some of them were successfully used as targets for protein structure elucidation (Redecke et al., 2013; Baskaran et al. 2015; Tsutsui et al. 2015; Nass et al., 2020; Lahey-Rudolph et al., 2021). Intracellular protein crystallization can occur naturally, providing distinct advantages for the cell (Schönherr et al., 2018), or as a consequence of heterologous gene expression in host cells. Depending on the target protein, crystallization efficiencies between more than 80 % and less than 1 % are observed (Lahey-Rudolph et al., 2020), and the intracellular crystal size may vary between the sub-micrometre range and several hundred micrometres (Schönherr et al., 2015; Schönherr et al., 2018). Targeting the crystals to different cell compartments or modifying the cells or the growth conditions may lead to optimized crystallization efficiency and/or larger crystals (Lahey-Rudolph et al., 2020; Mudogo et al., 2020), but identification of small crystals within a few cells of a large population may be required. Even if the resolution of light microscopy allows the crystal identification, this can be a time-consuming and laborious task. Thus, methods are required for the automatic characterization, screening and identification of intracellular crystals. Deep learning-based approaches can aid in this search.

Image segmentation is a method for partitioning an image into multiple disjoint regions via pixel-level classification. This is beneficial to locate objects and object boundaries. The technique is used in object detection and tracking (Maninis et al.,2018), with downstream applications like action recognition (Khan et al., 2021), autonomous driving (Kosecka et al., 1998), or scene understanding (Aarthi & Chitrakala, 2017). Image segmentation is increasingly used in medical image analysis as well, such as the macro and microscopic study of blood vessels (Ronneberger et al., 2015), tumor boundary detection (Havaei et al., 2017), or neuronal structures (Beier et al., 2017). Different approaches have been proposed for image segmentation (Nagaraja et al., 2015), based on Deep Neural Networks (DNN) (He et al., 2017; Chen et al., 2018), that produce high-quality results relying on enormous amounts of training data. Utilizing the transfer learning approaches (Pan & Yang, 2010), it is possible to bring a heavily, specifically trained model (for solving one problem) to solve a different (but related) problem. This is addressed by training the heavily trained model on few amount of data on this related problem.

In this work, we utilize the Mask R-CNN model (He et al.,2017), an extension of Faster R-CNN proposed by Ren et al. (Ren et al., 2015), adding object mask prediction. Faster R-CNN is a deep Convolutional Neural Network (CNN) for object detection which provides objectness score for each predicted object. The Mask R-CNN model generates (1) the bounding box around the object instances, (2) the objectness score, and (3) the segmentation of the object instance inside of the bounding box.

Two different types of *in cellulo* crystals have been used to establish the CNN-based instance segmentation method for protein crystal detection in living cells. The first target, HEX-1 from the fungus Neurospora crassa, referred here as ‘target H’, is a naturally self-assembling protein and main constituent of Woronin bodies in ascomycetes (Tenney et al., 2000). We recently reported that spontaneous self-assembly of HEX-1 into intracellular crystals is not restricted to the native environment of fungal cells (Lahey-Rudolph et al., 2020). On infection with a recombinant baculovirus encoding the HEX-1 gene, living insect cells also form regular, micrometre-sized hexagonal crystals that enabled structure elucidation using the fixed-target serial femtosecond crystallography approach (FT-SFX) at an XFEL (Lahey-Rudolph et al., 2021).

The second target, guanosine 5’-monophosphate reductase (GMPR) from the parasite Trypanosoma brucei, referred here as ‘target G’, catalyses the NADPH-dependent reductive deamination of guanosine 5’-monophosphate (GMP) to inosine 5’-monophosphate (IMP), a crucial step in the nucleotide metabolism of the parasite (Hedstrom 2012). GMPR has been structurally characterized by conventional X-ray crystallography (Imamura et al., 2020), but intracellular crystallization was not reported so far. However, this enzyme shows high structural homology to inosine 5’-monophosphate dehydrogenase (IMPDH) from T. brucei, the structure of which was recently elucidated by SFX using *in cellulo* crystals (Nass et al., 2020). Since both enzymes are part of the de novo purine biosynthesis cycle required for nucleotide production in parasitic protozoa, which differs significantly compared to that of mammals, these enzymes are considered as suitable antiparasitic drug targets (Bessho et al., 2016).

Using brightfield microscopy images from crystal containing insect cells we examine the performance of the Mask R-CNN algorithm, trained on different crystals, to identify different crystal shapes in the cellular environment, as a tool to extract information and aid in the development of protocols for the *in cellulo* crystallography field.

## 2. Material and Methods

### 2.1. Cloning and recombinant baculovirus production

The gene coding for guanosine 5’-monophosphate reductase (GMPR) from Trypanosoma brucei (GenBank Acc.No. XM839789.1) and the Woronin body major protein (HEX-1, GenBank Acc.No. XM_958614) from Neurospora crassa were amplified by PCR using primers 5’-CTAGGGTACCTC CTTCAATGAATCGGCATCC-3’ (sense) and 5’-CTAGGCTAGCAAGTTTGGCAACACCGTGAC-3’ (antisense), and 5’-CTAGGGTACCGGCTACTACGACGACGACG-3’ (sense) and 5’-CTAGGC TAGCGAGGCGGGAACCGTGG-3’ (antisense), respectively. The amplicons were cloned into pFastBac1 vector (Thermo Scientific) using KpnI and NheI restriction sites. The recombinant baculovirus (rBV) generation has previously been described (Lahey-Rudolph et al., 2020). In brief, recombinant bacmid DNA was generated by transformation of *E. coli* DH10EmBacY cells (Geneva Biotech) with the abovementioned vectors, purified with ZR Bac DNA Miniprep kit (Zymo Research) and used for lipofection of *Spodoptera frugiperda* Sf9 insect cells with Escort IV reagent (Sigma– Aldrich) according to the manufacturer’s instructions. The virus titre of the third-passage (P3) stock was calculated using the TCID_50_ (tissue-culture infectious dose; Reed & Muench, 1938) in a serial dilution assay as previously described (Lahey-Rudolph et al., 2020).

### 2.2. Cell culture, protein production and intracellular crystallization

High Five™ insect cells (Thermo Scientific) were grown at 27 °C and 100 rpm agitation in ESF921™ insect cell-culture medium (Expression Systems), keeping cell density between 0.4 - 2.5 × 10^6^ cells/ml. For intracellular crystallization of the target proteins, exponentially growing (1 × 10^6^ cells/ml) High Five cells were plated in 2 ml medium per six-well cell-culture plate and subsequently infected with the respective recombinant P3 baculovirus stock with a multiplicity of infection (MOI) of 0.1. After incubation at 27 °C for 44-72 h (GMPR) or 72-96 h (HEX-1) the cell cultures were imaged and *in cellulo* crystal formation was verified by light-microscopy.

### 2.3. Image acquisition parameters

Bright field images (2136×2136 px) were captured on a Nikon Qi-2 camera coupled onto a Nikon Ti2-E microscope mounting a Nikon S Plan Fluor ELWD 40x Ph2 ADM (NA=0.6) objective. The whole process was highly automatized, through random multipoint acquisition coupled to autofocusing (using Nikon’s Perfect Focus System) using the NIS Elements AR 5.41.01 software. Images were exported as 8 bit-RGB tifs and crystals were manually annotated using freeware Labelme (Russel et al., 2008) to produce the annotation of the crystals. Such annotations are used as ground-truth segmentations to train the Mask R-CNN model.

The Z sections of G-target or H-target containing cells (Figures 2 and 3) were taken using a Nikon AX-R confocal microscope, mounting a PlanApo λ 100X oil immersion objective (NA=1.45), using a pinhole size of 15.9 μm, a 488 laser for excitation and a YFP emission gate (518-551 nm) to detect the coexpressed cytosolic YFP. Similar optical Z-sections, of 1.05 μm and 1.20μm thickness, were obtained for target G and H, respectively.

### 2.4. Algorithm training and testing

The images on the train set of the two targets (H and G) are used to train and tune the pre-trained Mask R-CNN model. Basically, 50 images are annotated and used for training the model pre-trained on the COCO dataset (Lin et al., 2014).

The Mask R-CNN is a model for the object detection and segmentation. This model consists of a backbone of ResNet50 or ResNet100, the ResNet50 has less parameters and is faster in both train and test stages. We used the ResNet50 network for the bounding box recommendation of the model.

The model requires the images and the ground-truth masks of the same size as the input images to train the model weights. Therefore, we resized and converted the main images to the PNG format and the 512×512 px resolution. The trained model produces the bounding box around the detected crystals, score of the detection (the higher the better), and the binary segmentation of each of the crystals appearing inside the bounding boxes. The results of the model can be converted to images of any shape. This property of the model makes the possibility of training the model on smaller image resolutions depending on the computational power of the resources. Noting that training the model on larger image resolutions requires higher computational power. The model consists a region proposal network which produce the bounding boxes for the positions where the objects are found. Thereafter, the positions are given to a mask generation network to produce the pixel-wise segmentation of the objects inside the bounding boxes (Figure 1).

**Figure 1.**
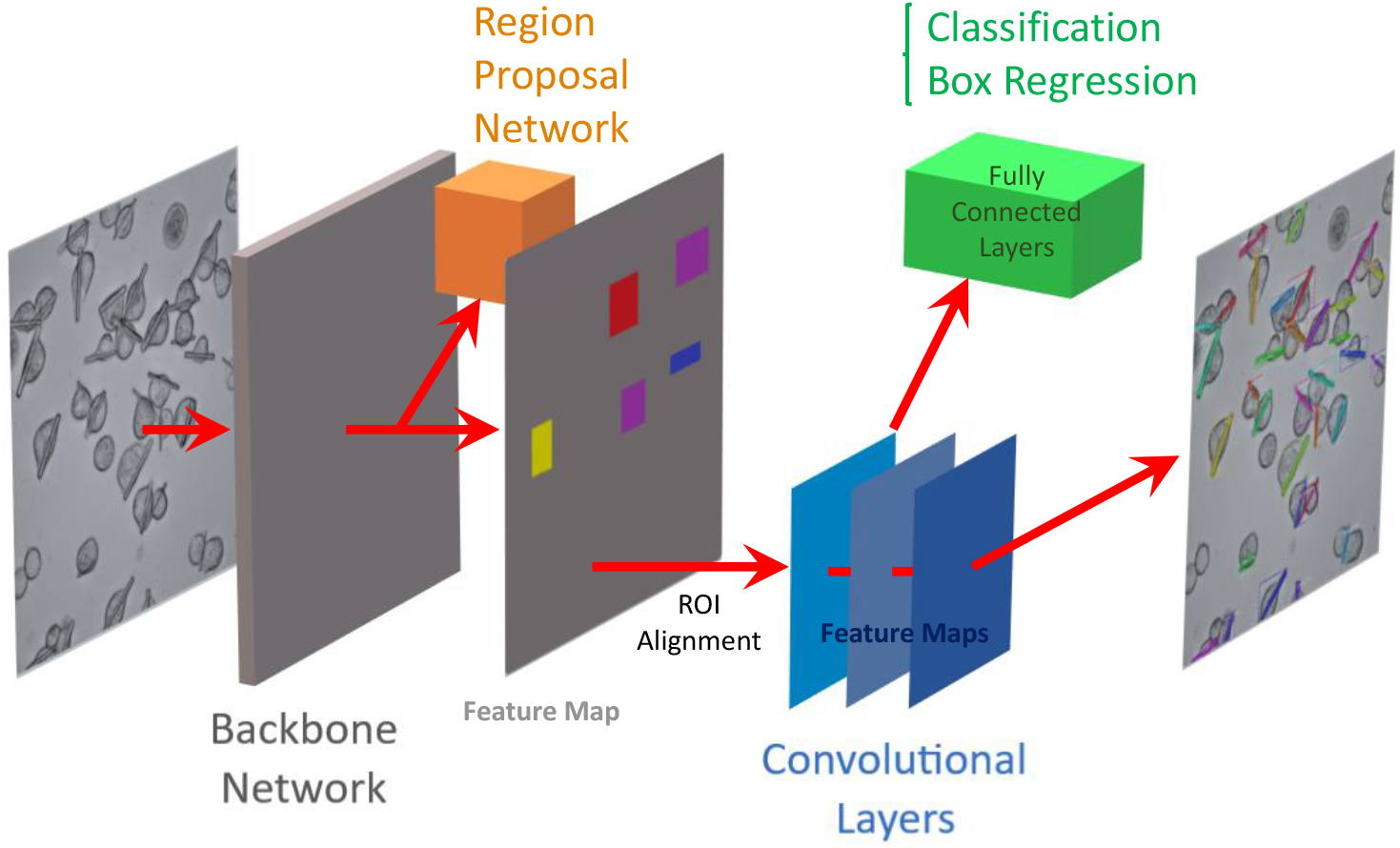
Schematic diagram showing the pipeline for training the pre-trained Mask R-CNN model (He et al., 2017) on our target G. Mask R-CNN adds a branch for object mask prediction, in parallel to bounding box recognition. The model generates bounding boxes, provides the objectness score for detection and accurately segments the intracellular crystals, requiring only few annotated images as training data set.

#### 2.4.1. Models Trained on Crystals of Type G and H

To segment the crystals of target G and H, separate models are trained on 50 images focusing on each type of the crystals for 40 epochs with the learning rate of 1e-3 and with the batch size of one via binary cross entropy loss function. To increase the robustness of the model, image augmentation is included such as random horizontal and vertical flip and cropping of the images. Therefore, two models are trained, one for target G and one for target H, where for both models the weights of the pre-trained Mask R-CNN model on COCO dataset are loaded. We used the Mask R-CNN implementation in [1]. Our method is similar to the approach of (Caicedo et al., 2019) for detection and segmentation of nuclei of cells in microscopy images. The trained Mask R-CNN on the data-science-bowl dataset (Caicedo et al., 2019) is used for detection and segmentation of the cells in our images (Figure 4).

We alternatively trained the pre-trained Mask R-CNN model on 10 and 30 images of each of the targets to study on the effect of the model by training on very few amount of training data. The training strategy is same as the one used with 50 train images, with the only difference being the number of images fed to the model for training.

#### 2.4.2. Incremental Learning on Targets G and H

To show the generalizability of the trained models on different targets we utilize the incremental learning approach (Geng et al., 2009). The model trained on one target is trained further on the target of different type. In this strategy, the trained models are tuned with 10 epochs and learning rate of 1e-3. To study on the effect of learning rate, the smaller value of 1e-4 is investigated.

## 3. Results and Discussion (style name: IUCr heading 1)

GMPR crystals started to grow after 44 h post-infection (p.i.), having a similar rectangular shape as observed for its close structural homolog IMPDH (Nass et al., 2020), as well and a comparable crystallization efficiency. At day 5 p.i., 95 % of the cells in the culture contained a single crystal, protruding from the cellular body, with its longer axis (that can reach up to 140 μm) orthogonal to the light path. The vast majority of the crystals tend to place at the bottom of the cells, as observed with confocal optical sectioning (Figure 2). HEX-1 crystals have been observed in the baculovirus-infected cells at day 4 p.i., exhibiting the reported hexagonal shape and the expected crystallization efficiency (>85 %), without protruding the cell body (Lahey-Rudolph et al, 2021). Next to the smaller size, HEX-1 crystals show a significant degree of clustering and grow in diverse orientations (Figure 3), adding a level of complexity to crystal analysis compared to GMPR, for both manual annotation and automatic segmentation.

**Figure 2.**
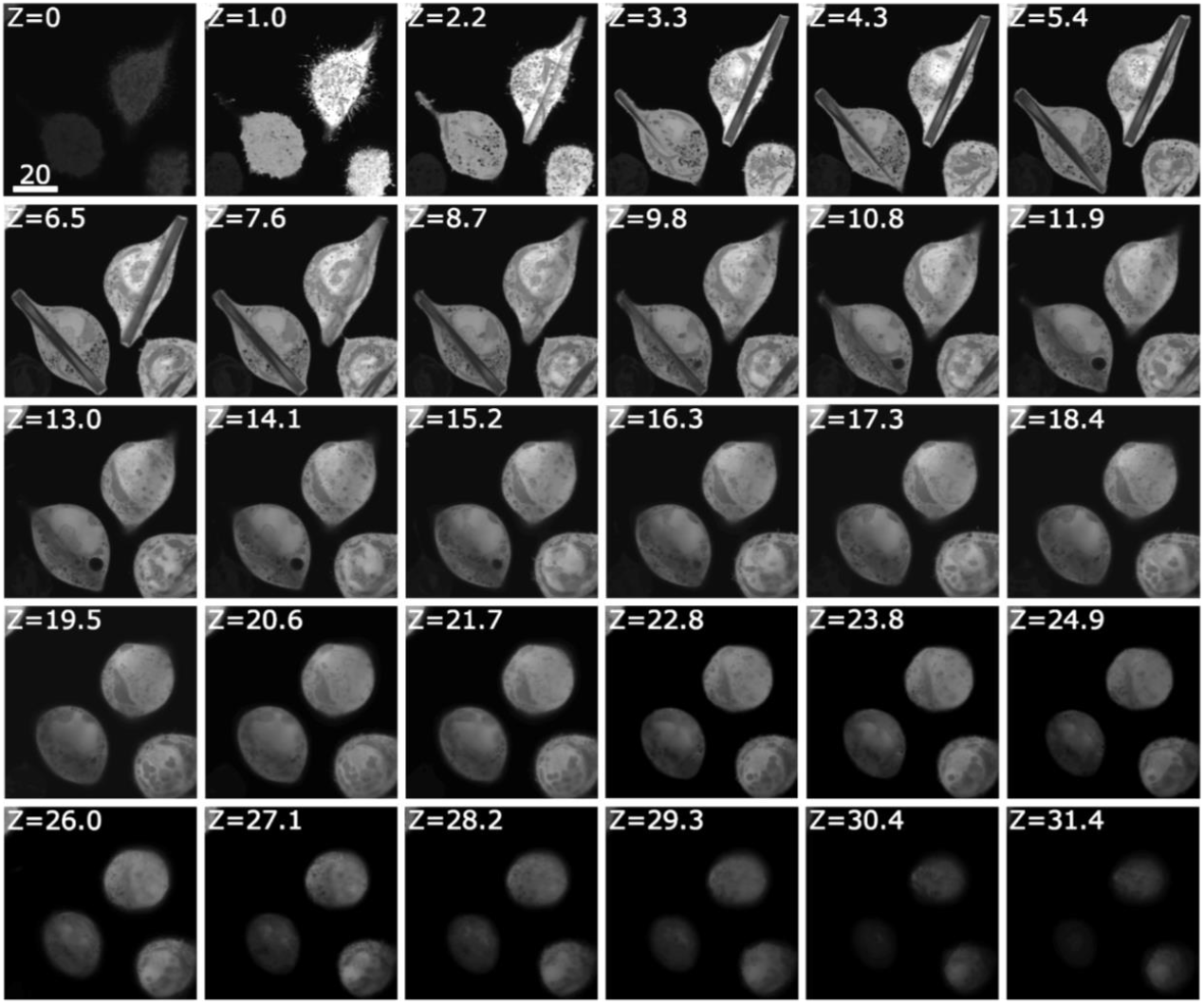
Fluorescence confocal optical sectioning. Fluorescence signal is produced by the coexpressed cytosolic YFP, while G-target (GMPR) crystals are identified by the absence of signal. Z value of 0 corresponds to the bottom of the cell. The optical sectioning (1.05 μm steps) allows to confirm the horizontal position of the crystals at the cell base. All values correspond to μm units.

**Figure 3.**
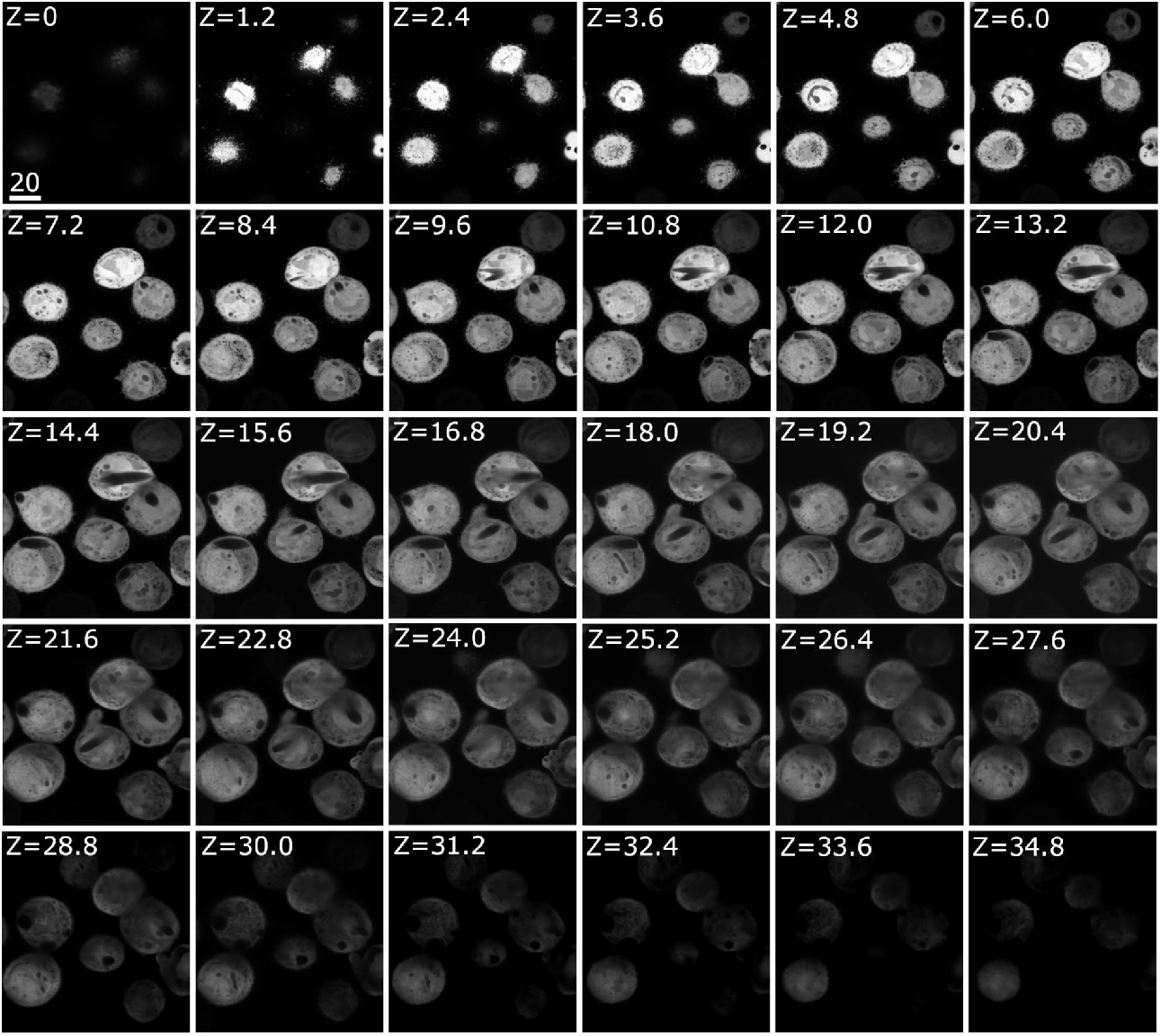
Fluorescence confocal optical sectioning. Fluorescence signal is produced by the coexpressed cytosolic YFP, while H-target (Hex-1) crystals are identified by the absence of signal. Z value of 0 corresponds to the bottom of the cell. The optical sectioning (1.20 μm steps) confirms the relatively random orientation and position of the crystals throughout the cell body, as well as the presence of clustered, multishaped crystals. All values correspond to μm units.

The first parameter to analyze is the impact of the number of images to train the Mask R-CNN algorithm, for each respective target (Table1, rows 1-6; Figures S1-S3). For each scenario, the performance of the trained models is evaluated on test sets of two targets G and H, each containing 150 images. We calculated the performance on the basis of three indicators: F-measure, Jaccard-index, and Δ object. Concretely, F-measure shows the contour matching score between the predictions and the annotations, Jaccard Index is used to estimate shape-matching accuracy, and Δ object indicates the number of unrecognized objects. For the first two indicators, higher values indicate a better algorithm performance, whereas for Δ object, the lower the value, the better. The results show that just 10 images were sufficient to train the algorithm to recognize each of the targets, with just a slight increase in performance by the addition of more images (up to 50) in the case of target G (Table 1, rows 1-6; Figures S1-S3). Due to that, we chose 50 images as the best compromise, thus selecting this as a base for the following training strategies. Regarding cell recognition, the model trained with the science-bowl dataset is able to recognize the cell bodies (Figure 4) based on previous training approaches (Mela & Liu, 2021). Our results indicate that learning approaches are useful for recognizing *in cellulo* crystals, using a limited number of epochs and annotated images.

**Table 1.** Validation of the model. The quantitative results are reported based on F-measure, Jaccard index, and Δ object. On the Δ object, the lower this value the better, whereas higher values for F-measure and Jaccard index indicates a better performance. The values are reported on the indicated test sets (target H or G).

|  | *Avg. $\Delta$ object* | *Avg. F-measure (%)* | *Avg. Jaccard Index (%)* |
| --- | --- | --- | --- |
|  | *target H/target G* | *target H/target G* | *target H/target G* |
| *TS10G* | 20.62/6.01 | 47.14/75.67 | 31.41/60.94 |
| *TS30G* | 16.49/5.38 | 58.95/77.11 | 42.23/62.82 |
| *TS50G* | 18.69/5.29 | 54.95/77.70 | 38.38/63.60 |
| *TS10H* | 5.15/7.37 | 82.19/54.40 | 69.92/37.65 |
| *TS30H* | 4.37/12.89 | 81.61/55.07 | 69.10/38.36 |
| *TS50H* | 3.60/15.15 | 82.51/57.77 | 70.38/40.97 |
| *TS50G10H (LR=0.001)* | 5.15/4.43 | 82.35/75.34 | 70.13/60.52 |
| *TS50G10H (LR=0.0001)* | 5.64/3.61 | 78.79/77.23 | 65.18/62.97 |
| *TS50G30H (LR=0.001)* | 4.58/5.28 | 81.76/76.20 | 69.27/61.63 |
| *TS50G30H (LR=0.0001)* | 5.46/3.32 | 79.92/77.34 | 66.69/63.12 |
| *TS50G50H (LR=0.001)* | 4.27/7.19 | 83.34/75.64 | 71.55/60.90 |
| *TS50G50H (LR=0.0001)* | 5.44/4.04 | 80.93/76.89 | 68.11/62.52 |
| *TS50H10G (LR=0.001)* | 15.29/6.55 | 63.88/73.77 | 47.41/58.53 |
| *TS50H10G (LR=0.0001)* | 9.63/3.68 | 73.49/62.75 | 58.36/45.85 |
| *TS50H30G (LR=0.001)* | 11.79/4.68 | 67.49/73.01 | 51.26/57.61 |
| *TS50H30G (LR=0.0001)* | 9.61/4.43 | 73.65/60.29 | 58.57/43.30 |
| *TS50H50G (LR= 0.001)* | 13.82/6.55 | 68.36/74.26 | 52.33/59.16 |
| *TS50H50G (LR= 0.0001)* | 9.39/4.22 | 73.47/59.93 | 58.38/42.94 |

**Figure 4.**
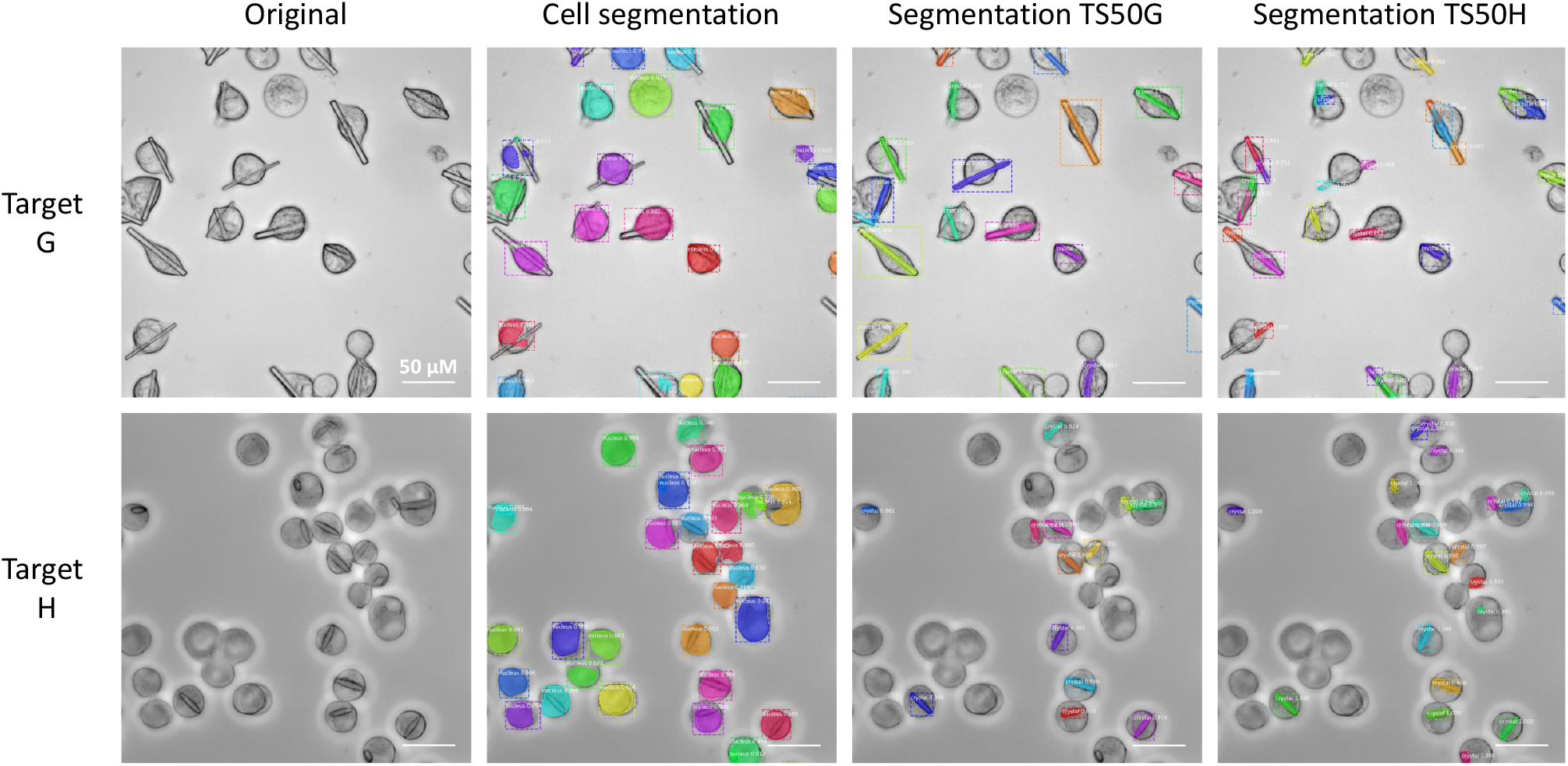
Primary training of the algorithm on 50 images of target G (TS50G) or target H (TS50H) and evaluation on each of the targets (indicated on the left side of the panel). A trained algorithm on the data-science-bowl dataset (Caicedo et al.,2019) was used for segmentation of the cell bodies. The results indicate that specific training on the target crystal is required for optimum performance.

We next investigated the ability of the trained algorithm to recognize crystal targets of different nature than that of the ones used for generating the training set. For doing that, the previously trained algorithms (on H or G targets) were used to recognize the other (G or H, respectively) target crystals. Despite the trained algorithms were able to recognize some of the “new” target features, the quality of the predictions was lower in both cases, as expected (Table 1, rows 1-6; Figure 4, S1-S3).

In light of these results, we tried to follow the incremental learning approach, based on the continuation of the previous training (that was “off target”) by adding few (10) annotated images from the target to be tested. In this approach, we set two different learning rates (0.001 and 0.0001) to test the influence of this parameter in the outcome.

The results (Table1, rows 7-18; Figure 5; Figures S4-S6) indicate that the addition of these few images (of the target to be tested) to the training process (referred here as “secondary” training) improved the recognition of this “secondary” target significantly, and for both targets. However, when looking the performance of the resulting algorithms on their “primary” targets, the effect of this “secondary” training substantially differs. Thus, in the case of the algorithm “primarily” trained on target H (smaller, more complex crystals), the “secondary” training (providing “versatility” towards target G) comes across with a negative impact on the recognition of its “primary” target, as observed when comparing the indicator values with that of the algorithm trained exclusively for that target (Table 1; Figures S4-S6). That finding suggests that, in some cases, widening the scope of targets to be recognized is at expense of sacrificing specific performance. On the other hand, this behaviour is not observed when G (larger, less complex crystals) was the target used for the primary training, suggesting that the approach works optimally when the primary training is done with simpler cases, but not the other way around. In the latter cases, the negative impact is more evident when using higher learning rates for the “secondary” training, with half of the negative impact observed when using lower learning rates. Interestingly, this negative impact, inversely correlates with the benefits seen on the recognition of the “secondary” target, which are higher when using higher learning rates. A progressive increase of the images used for this “secondary” training did not result in a clear change in performance, suggesting that 10 images are sufficient to generate the observed changes.

**Figure 5.**
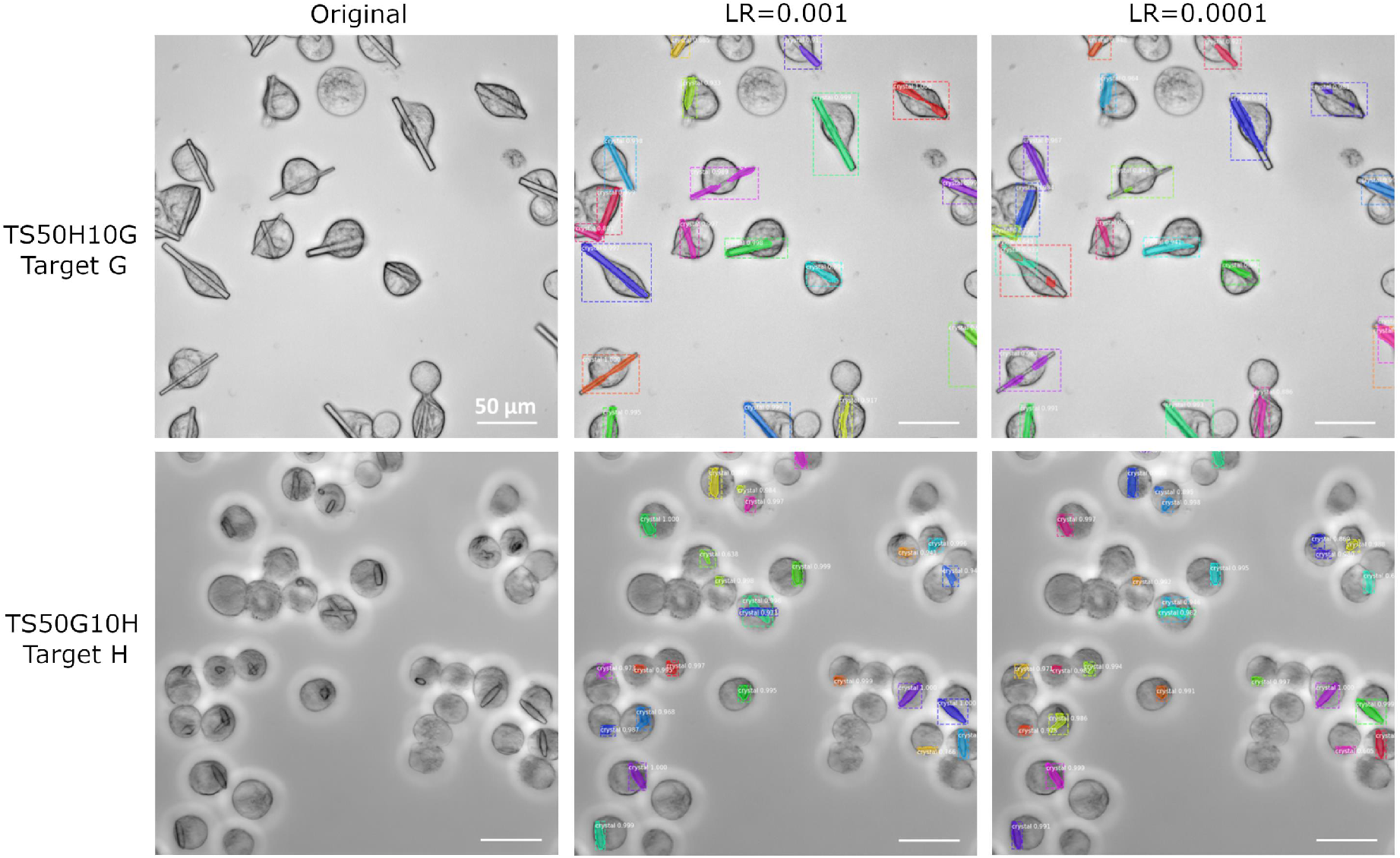
Secondary training of TS50H algorithm with 10 images of target G (TS50H10G, upper row), and TS50G algorithm with 10 images of target H (TS50G10H, lower row), using different learning rates, and evaluated on G or H targets, respectively. For this secondary training, the higher learning rate (0.001) provides better results in the case of G target, whereas lower learning rates works better in the case of target H. That shows that the learning rate is a parameter that must be tuned depending on the target type.

It is important to note that the secondary training does not seem to increase, for any of the two crystals, the performance of the algorithm on the recognition of its primary target. That demonstrates, on one hand, that no additional information is fetched from a different crystal type, and on the other hand, indicates that the algorithm is very efficient in extracting all the necessary information from very few images.

Altogether, the results confirm the effortless adaptability of the algorithm, and the potential to improve it with every new target on study. However, this incremental learning approach, in some cases, may interfere with the previous knowledge of the algorithm, making it to “unlearn”, improving the versatility while reducing specificity. The current manuscript sheds some light into this behaviour, and indicates that the approach requires to be tuned depending on the shape characteristics of the objects used for training the algorithm.

## 4. Conclusion

We showed the potential of our method for the segmentation of *in cellulo* crystals. Concretely, we propose a model based on Mask R-CNN, which accurately detects different types of intracellular crystals, harbouring well differentiated shapes. The model can be further tuned and effortlessly adapted to new crystal shapes. In addition, optical sectioning allows to segment the cells and crystals in different layers, opening the scope to 3-dimensional segmentation of the objects.

The current set-up, including fully automated acquisition, aims to be used as a general screening pipeline to rapidly score cell cultures for successful intracellular crystal growth. It is particularly useful for cells producing unknown protein targets with low or very low crystallization efficiency, preventing hours of manual cell scoring. The algorithm can also assist in the selection of optimization protocols during *in cellulo* crystallization, being able to monitor the impact of the different approaches on the occurrence or the size or shape of the crystals. Potentially, it can be used to perform real-time tracking of cells containing crystals (or isolated crystals) during X-ray diffraction experiments at synchrotrons or FELs, either placed on fixed targets or flowing through high viscosity jets. That could result in significant advantages, like avoiding the need to irradiate the sample for target localization, or synchronizing beam exposures to selected targets or crystal shapes.

## 5. Acknowledgements

This research was supported in part through the Maxwell computational resources operated at Deutsches Elektronen-Synchrotron DESY, Hamburg, Germany. This work is in part supported by funding from the German Federal Ministry for Education and Research (BMBF; grant 05K18FLA). The authors thank Sophie Nachtschatt, University of Lübeck, Germany, for cloning the HEX-1 and GMPR genes into the pFastBac vectors and for bacmid generation, and Roman Weissblatt, European XFEL GmbH, Germany, for helping with manual image annotation.

