## Supporting Information for "Convolutional neural network approach for the automated identification of *in cellulo* crystals"

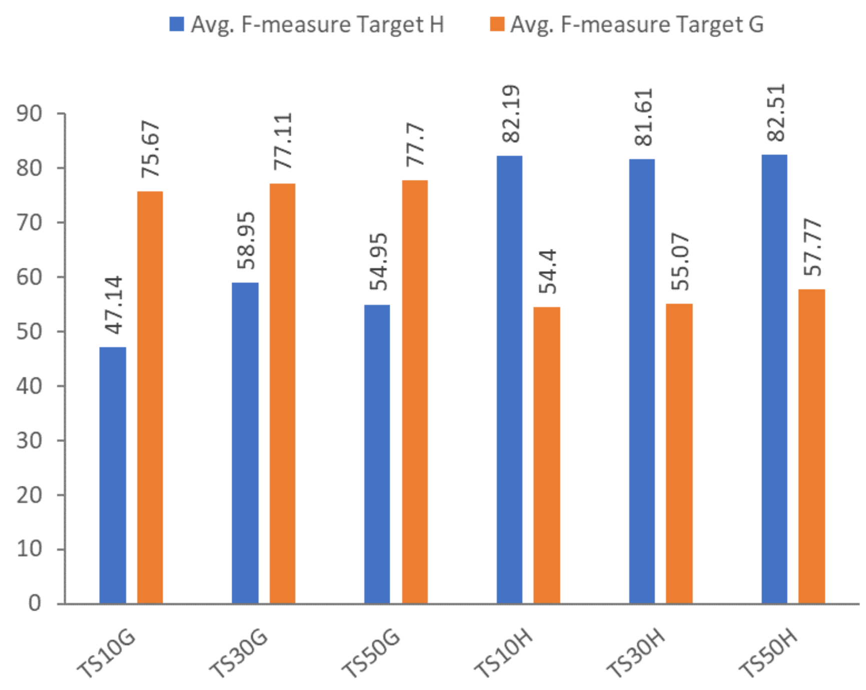


**Figure S1** Average F-measure Index showing the effect of the primary training on the recognition of each of the targets. Graphical representation of the values of the F-measure Index indicated in Table 1, corresponding to the different training strategies (X axis labels), tested on each of the targets, H or G. F-measure determines the contour matching score between the predictions and the annotations, so higher values correspond to better performance.

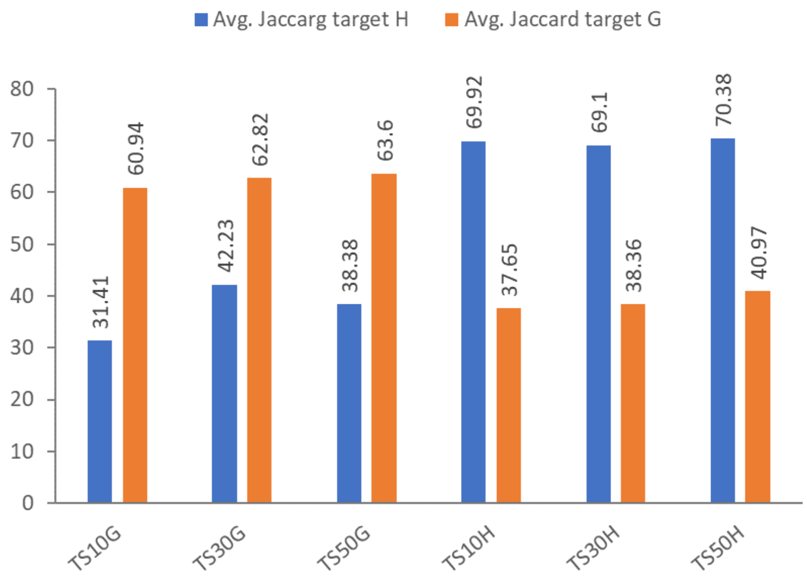


**Figure S2** Average Jaccard Index showing the effect of the primary training on the recognition of each of the targets. Graphical representation of the values of the Jaccard Index indicated in Table 1, corresponding to the different training strategies (X axis labels), tested on each of the targets, H or G. Jaccard Index allows to estimate shape-matching accuracy, so higher values correspond to better performance.

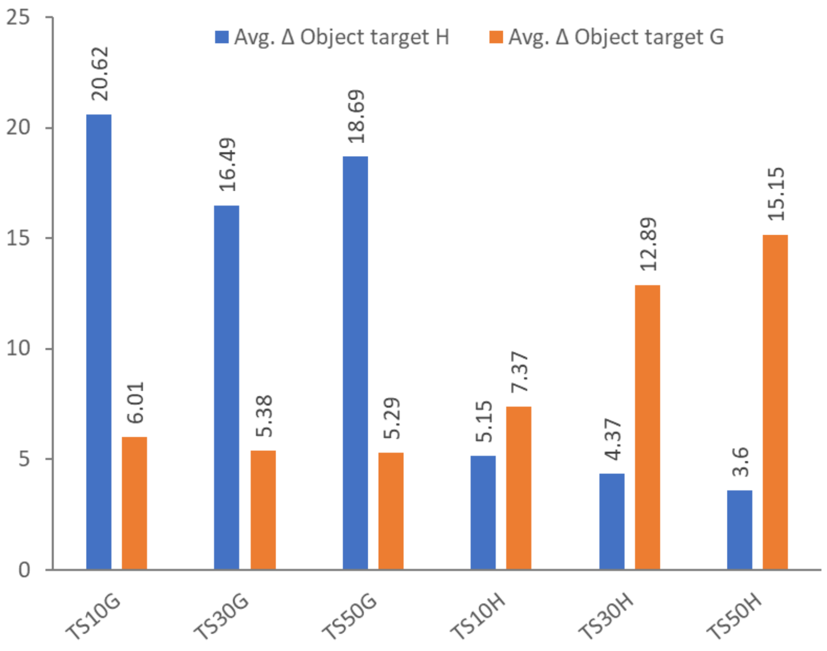


**Figure S3** Average Δ Object showing the effect of the primary training on the recognition of each of the targets. Graphical representation of the values of the Δ Object indicated in Table 1, corresponding to the different training strategies (X axis labels), tested on each of the targets, H or G. Δ Object is an indicator of the number of unrecognized objects, so lower values correspond to better performance.

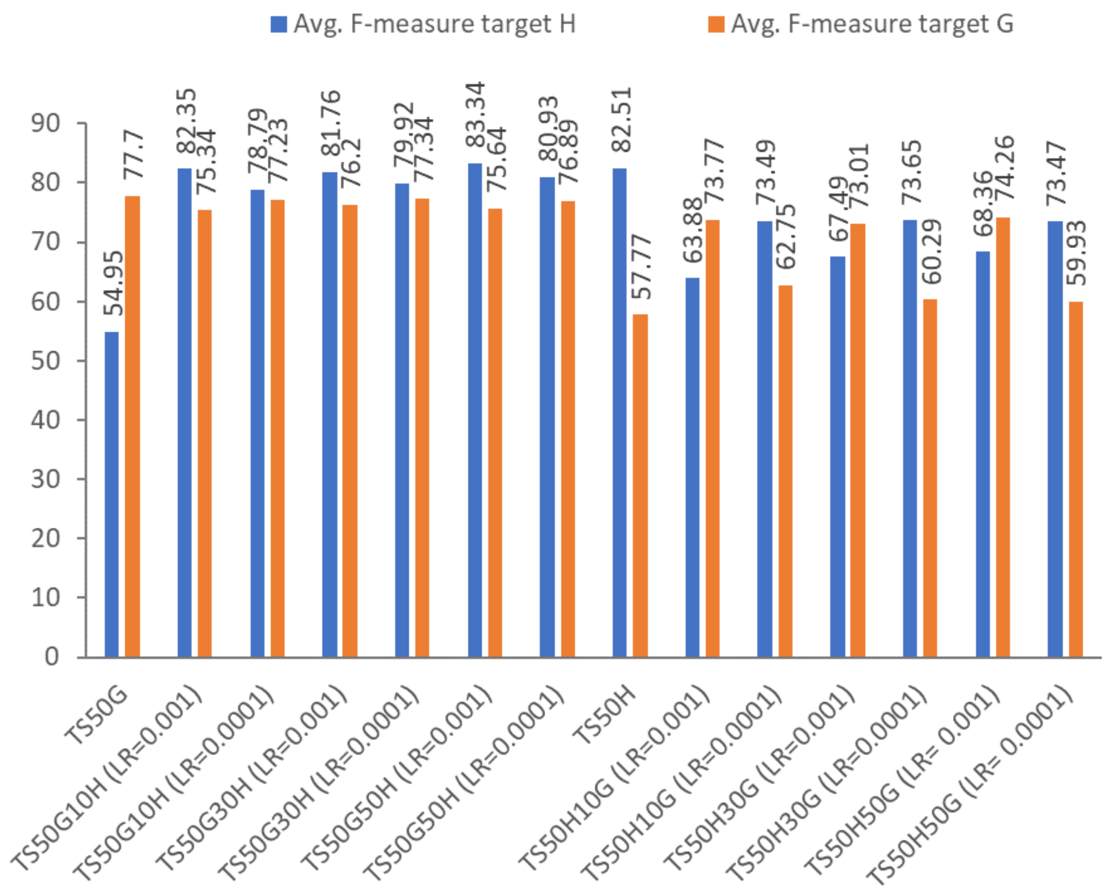


**Figure S4** Average F-measure Index showing the effect of the secondary training on the recognition of each of the targets. Graphical representation of the values of the F-measure Index indicated in Table 1, corresponding to the different training strategies (X axis labels), tested on each of the targets, H or G. F-measure determines the contour matching score between the predictions and the annotations, so higher values correspond to better performance.

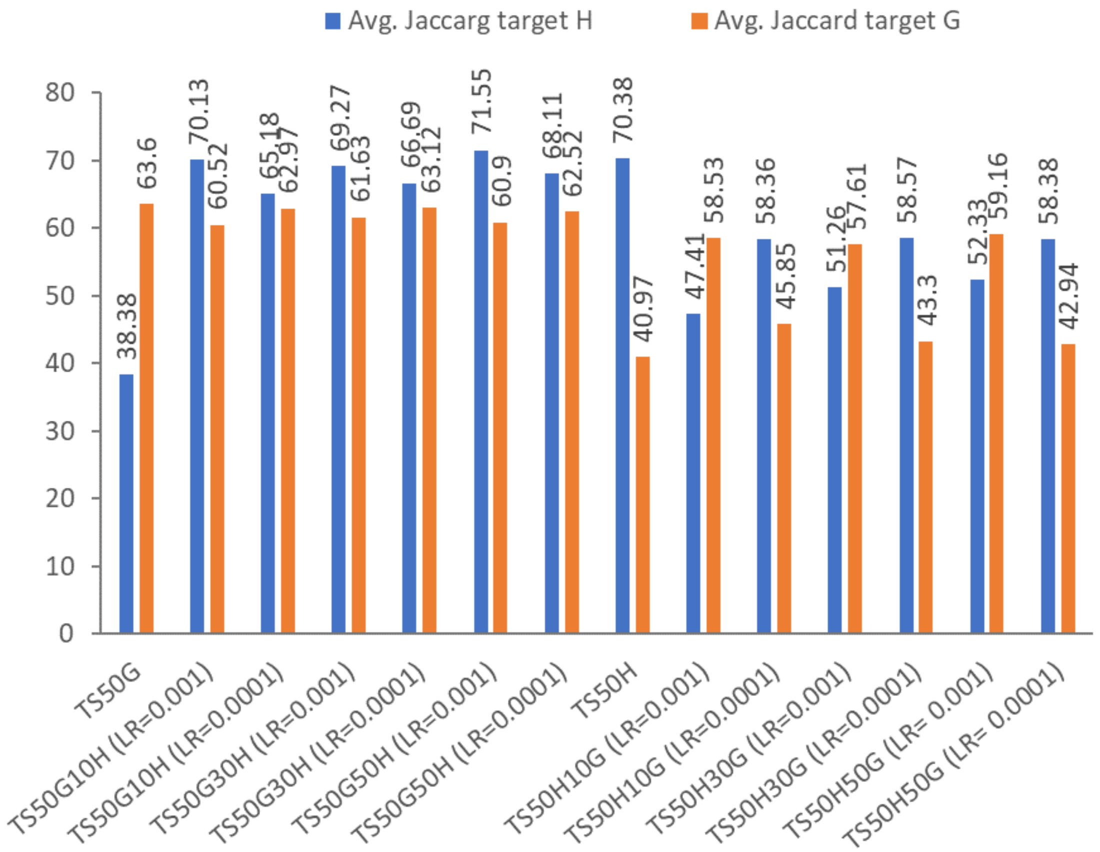


**Figure S5** Average Jaccard Index showing the effect of the secondary training on the recognition of each of the targets. Graphical representation of the values of the Jaccard Index indicated in Table 1, corresponding to the different training strategies (X axis labels), tested on each of the targets, H or G. Jaccard Index allows to estimate shape-matching accuracy, so higher values correspond to better performance.

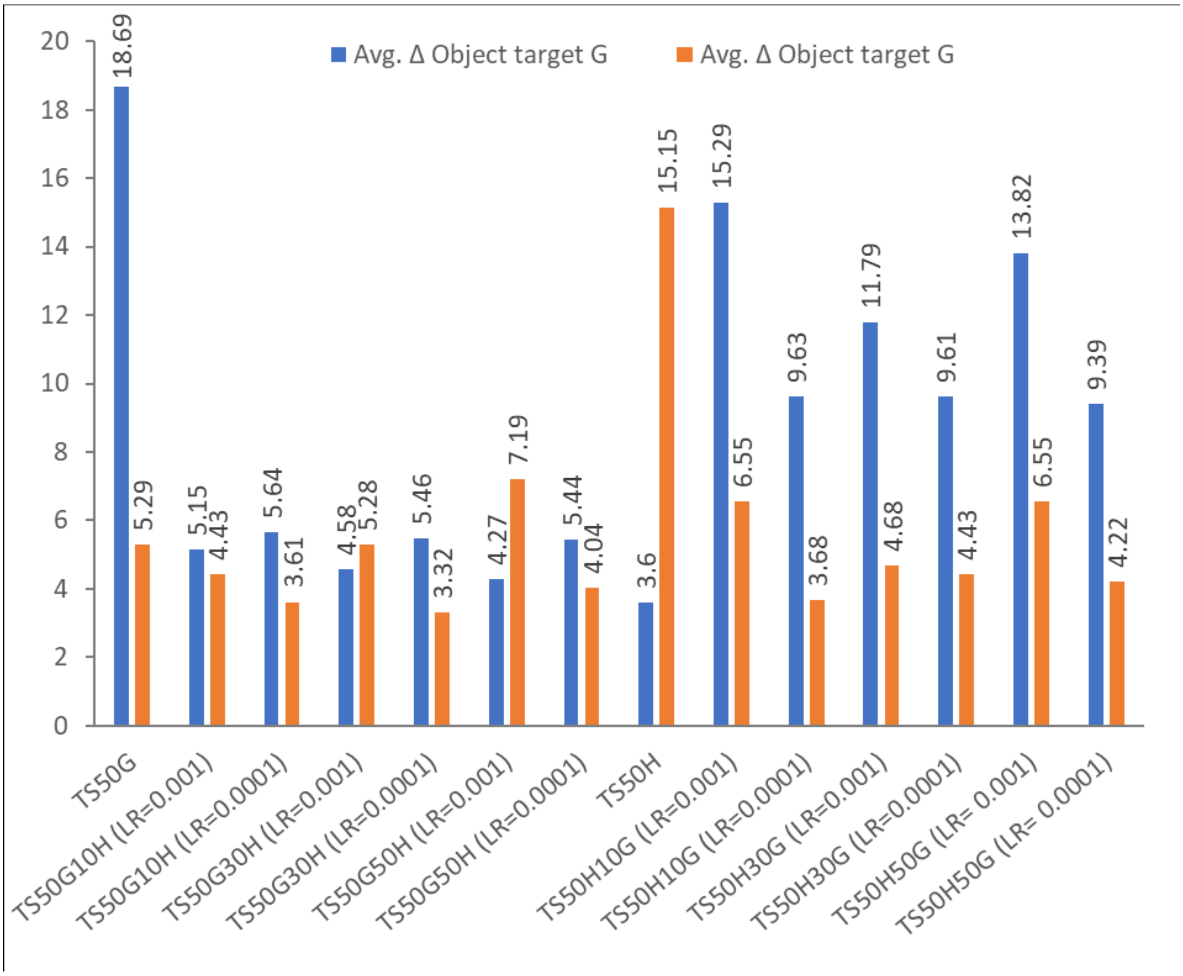


**Figure S6** Average Δ Object showing the effect of the secondary training on the recognition of each of the targets. Graphical representation of the values of the Δ Object indicated in Table 1, corresponding to the different training strategies (X axis labels), tested on each of the targets, H or G. Δ Object is an indicator of the number of unrecognized objects, so lower values correspond to better performance.
